# Human Dermal Fibroblast Subpopulations and Epithelial Mesenchymal Transition Signals in Hidradenitis Suppurativa Tunnels are Normalized by Spleen Tyrosine Kinase Antagonism in Vivo

**DOI:** 10.1101/2023.02.23.529664

**Authors:** A Flora, R Jepsen, EK Kozera, JA Woods, GD Cains, M Radzieta, SO Jensen, M Malone, JW Frew

**Affiliations:** Laboratory of Translational Cutaneous Medicine, Ingham Institute for Applied Medical Research, Sydney, Australia; University of New South Wales, Sydney, Australia; Department of Dermatology, Liverpool Hospital, Sydney, Australia; Holdsworth House Medical Practice, Sydney, Australia; South West Sydney Limb Preservation and Wound Research. Ingham Institute for Applied Medical Research; School of Medicine, Western Sydney University, Sydney, Australia

## Abstract

Hidradenitis Suppurativa is a chronic inflammatory disease of which the pathogenesis is incompletely understood. Dermal fibroblasts have been previously identified as a major source of inflammatory cytokines, however information pertaining to the characteristics of subpopulations of fibroblasts in HS remains unexplored. Using in silico-deconvolution of whole-tissue RNAseq, Nanostring gene expression panels and confirmatory immunohistochemistry we identified fibroblast subpopulations in HS tissue and their relationship to disease severity and lesion morphology. Gene signatures of SFRP2+ fibroblast subsets were increased in lesional tissue, with gene signatures of SFRP1+ fibroblast subsets decreased. SFRP2+ and CXCL12+ fibroblast numbers, measured by IHC, were increased in HS tissue, with greater numbers associated with epithelialized tunnels and Hurley Stage 3 disease. Pro-inflammatory CXCL12+ fibroblasts were also increased, with reductions in SFRP1+ fibroblasts compared to healthy controls. Evidence of Epithelial Mesenchymal Transition was seen via altered gene expression of SNAI2 and altered protein expression of ZEB1, TWIST1, Snail/Slug, E-Cadherin and N-Cadherin in HS lesional tissue. The greatest dysregulation of EMT associated proteins was seen in biopsies containing epithelialized tunnels. The use of the oral Spleen tyrosine Kinase inhibitor Fostamatinib significantly reduced expression of genes associated with chronic inflammation, fibroblast proliferation and migration suggesting a potential role for targeting fibroblast activity in HS.

## INTRODUCTION

Hidradenitis Suppurativa (HS) is a chronic inflammatory skin disease with an estimated prevalence of 1% globally (Sabat et al., 2020; Wolk et al., 2020). Lesions include nodules and abscesses as well as epithelialized dermal tunnels and significant hypertrophic scarring and contractures in flexural areas (Frew et al., 2021a). The molecular pathogenesis of HS is incompletely defined, but dermal fibroblasts are known to be major contributors of proinflammatory cytokines including IL-1, CCL20, CXCL1, CXCL8, IL36B and SERPINB4 (Witte-Handel et al., 2018; Zouboulis et al., 2020). Fibroblasts are important drivers of wound healing and epidermal repair (Artuc et al., 2002). The mechanisms of wound healing involve activation and migration of fibroblasts as well as epithelial-mesenchymal transition (EMT) like mechanisms which are proposed to be dysregulated in HS. (Sen and Roy., 2021; Artuc et al., 2002) It has been hypothesized that fibroblasts may play a role in the development of epithelialized tunnels through keratinocyte-fibroblast interactions and EMT. (Frew et al., 2019a). The evidence for activation of EMT in lesions of HS is limited and further exploration of this hypothesis may lead to novel therapeutic targets against the progression of disease (Nelson et al., 2019; Frew et al., 2021b) including the development of tunnels, hypertrophic scarring and flexural contractures.

Dermal fibroblasts are a heterogeneous population, with functionally distinct fibroblast subpopulations have been identified in normal skin (Ascencion et al 2021; Deng et al., 2021). More recent investigations have identified fibroblast subpopulations in chronic inflammatory dermatoses including scleroderma (Deng et al., 2021), and cutaneous lupus (Mizoguchi et al., 2018). Distinct fibroblast subpopulations demonstrate different effector functions which may have differing contributions to cutaneous inflammation and scarring (Ascenscion et al., 2021; Tabib et al., 2018; Tabib et al., 2021). The contribution of fibroblasts to the pathogenesis of HS has been proposed (Frew et al., 2019a), but HS-specific investigations have been limited to date. (Frew et al., 2021b).

The therapeutic modulation of fibroblasts is an emerging area of research in fibroinflammatory diseases with existing therapies including Janus Kinase (JAK) inhibitors and Spleen Tyrosine Kinase (SYK) inhibitors demonstrating evidence of clinically relevant fibroblast modulation (Liu et al., 2019). These drugs have the potential to be repurposed for investigation into their benefit in the setting of HS. SYK inhibition has the additional benefit of potentially addressing B cells, plasma cells and monocytes which are significantly dysregulated in HS (Gudjonsson et al., 2020, Musilova et al., 2020, Byrd et al., 2019; Frew et al 2020).

In this study we aimed to examine the presence of fibroblast subpopulations in lesional tissue of HS compared to non lesional tissue and healthy controls. Additionally, the effect of SYK antagonism with Fostamatinib upon gene expression profiles in lesional tissue was assessed.

## RESULTS

### The Fibroblast Transcriptome is Upregulated in Lesional vs non-Lesional Tissue of Hidradenitis Suppurativa

Whole tissue RNAseq from lesional and non-lesional tissue from 20 individuals with HS identified differentially expressed genes associated with fibroblast activity including *TGFB1, OSM, IL6, MMP1* and *MMP3*. These genes discretely clustered in lesional versus non lesional tissue (Figure 1a). This demonstrates concordance with other published RNAseq datasets (Hoffman et al., 2018; Zouboulis et al., 2020). Fibroblast associated genes such T*HY1, LUM, FN1* and *COL1A1* were significantly overexpressed in lesional compared with non lesional HS tissue (Figure 1b, Supplementary Data). Examining genes associated with identified fibroblast subpopulations (Tabib et al., 2018; Tabib et al., 2021; Ascencion et al., 2021) did identify differentially expressed genes in whole tissue RNAseq including *SFRP1, SFRP2* and *DPP4*. Genes associated with Epithelial Mesenchymal Transition (EMT) including *CDH1, CDH2, SNAI2, TWIST1, ZEB1* were not significantly differentially expressed between lesional and non-lesional HS tissue (Supplementary Data). Pathway analysis demonstrated positive enrichment of pathways associated with tissue remodeling, extracellular matrix disassembly and collagen fibril organization. (Figure 1b).

**Figure 1:**
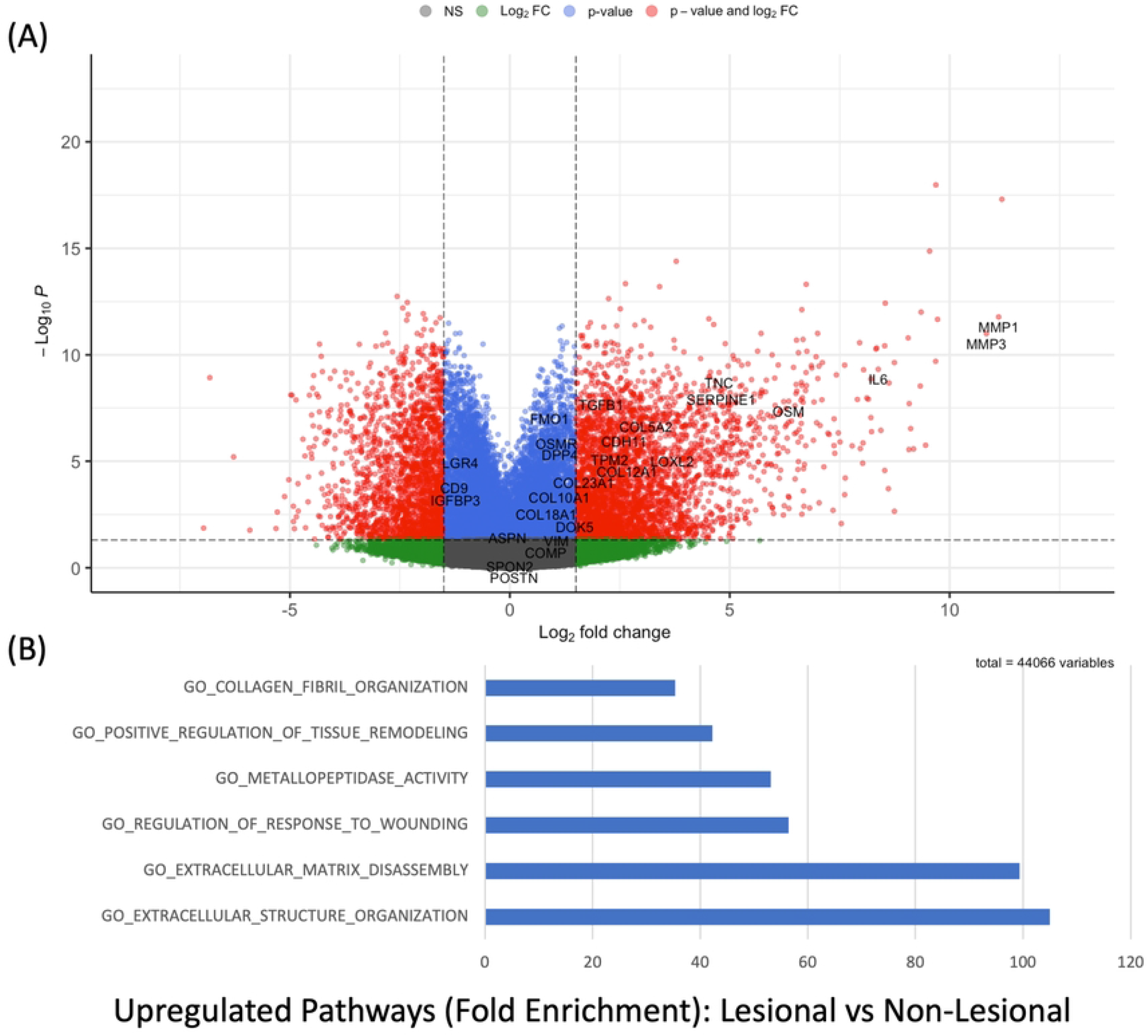
(A) Volcano plot of differentially expressed genes in lesional versus non-lesional tissue in Hidradenitis Suppurativa. Genes pertaining to fibroblasts and fibroblast associated gene expression are highlighted including MMP (Matrix metalloproteinase) 1 and 3, Oncostatin M (OSM), transforming growth factor Beta 1 (TGFB1), vimentin (VIM) and Dipeptidyl protease 4 (DPP4). (B): Pathway analysis by PANTHER demonstrating enrichment of pathways pertaining to tissue remodeling, metalloproteinase activity and other extracellular matgric and fibroblast associated functions in lesional, versus non lesional tissue in HS.

### In Silico Deconvolution of the HS Tissue Transcriptome Identifies Differential Expansion of Fibroblast Subpopulations in Lesional vs Non-Lesional Tissue

Using CibersortX, in silico deconvolution of whole tissue RNAseq was used to identify transcriptomic signatures indicative of fibroblast subpopulations (as defined by Ascencion et al., 2021) (Supplementary Figure 2). No significant difference between lesional and non-lesional tissue was identified in the cell proportions of the overall fibroblast population (Figure 2a). However, the differential cell proportion of fibroblast sub populations (Ascension et al., 2021) was significantly between lesional and non-lesional tissue (Figure 2b-e). SFRP2+ fibroblast (Type A) gene signatures were upregulated in lesional tissue, with a reduction in SFRP1+ gene signature (Type C) in lesional tissue and no significant change between CXCL12+ (Type B) gene signatures in lesional tissue compared to non-lesional tissue (Figure 2c-e).

**Figure 2:**
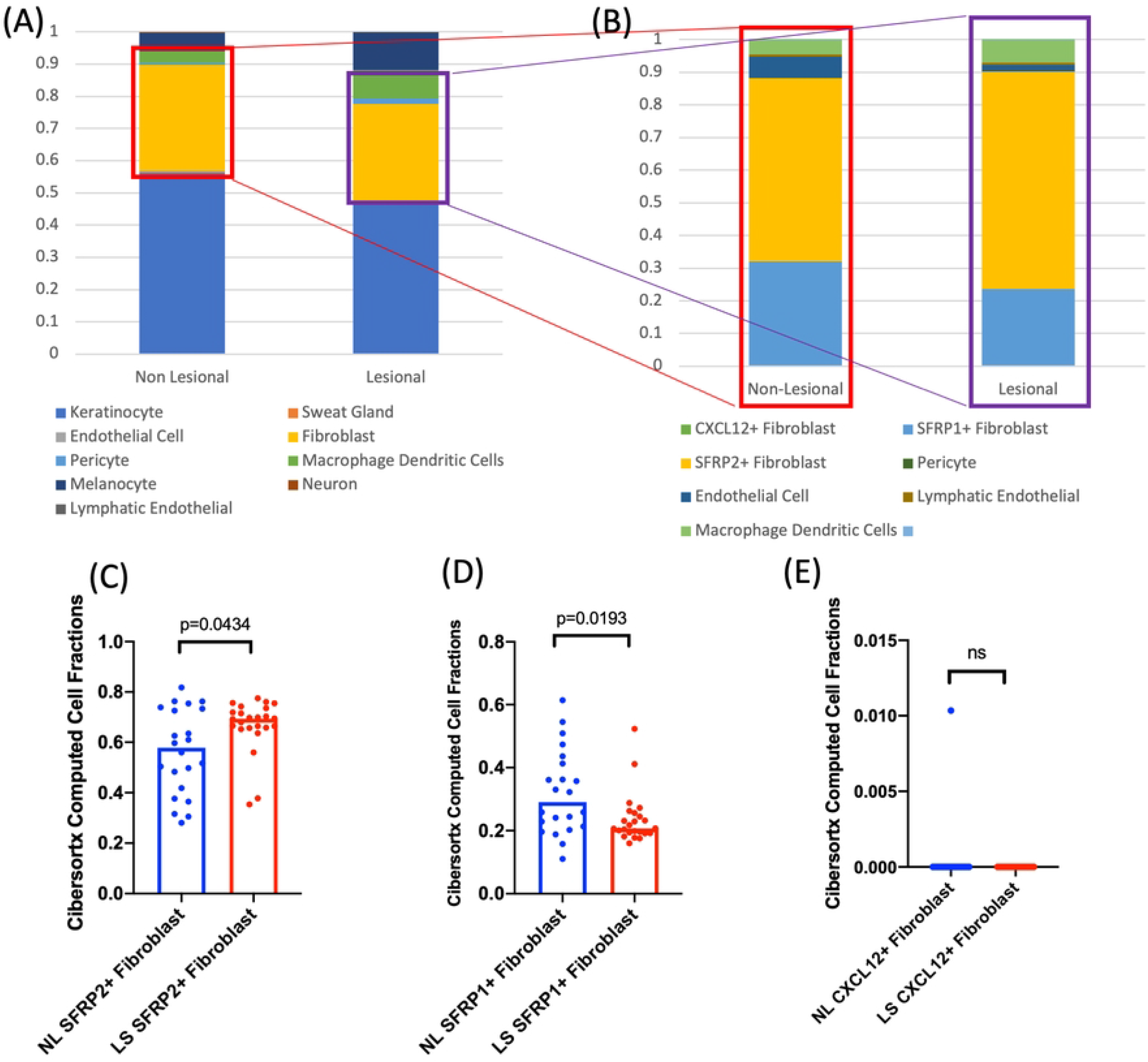
In-Silico Deconvolution of whole tissue RNAseq demonstrates differential expansion of fibroblast subpopulations in lesional vs non lesional tissue of Hidradenitis Suppurativa. (A): Cell Proportion data as calculated by CIbersortX in-silico deconvolution demonstrating alterations in cell populations between lesional and non lesional tissue. (B) Isolation of only fibroblast subsets indicates an expansion of SFRP2+ fibroblast subpopulartion gene signautres. (C): Individual sample data identifies a statistically signficiant expansion of SFRP2+ fibroblast subsets in lesional tissue, decreased SFRP1+ populations and no significant change in CXCL12+ populations in whole tissue RNAseq with in-silico deconvolution.

### Immunohistochemistry validates the presence of discrete fibroblast subpopulation locations in Hidradenitis Suppurativa tissue

Transcriptomic findings illustrating differential gene expression of fibroblast sub-populations was validated using immunohistochemistry with primary antibodies against SFRP1, SFRP2 and CXCL12 proteins (primary antibodies used listed in Supplementary Data) in lesional and non lesional HS tissue (Figure 3a-c). Significant differences in the number of SFRP1+, SFRP2+ and CXCL12+ cells (measured using semiquantitative IHC staining) was seen between normal and HS lesional tissue (Figure 3d-f). Stratification by Hurley stage illustrated a significant reduction in SFRP1+ cells and an elevation in the number of SFRP2+ and CXCL12+ cells in Hurley Stage 3 patients compared with Hurley stage 2 patients (Figure 3d-f). Stratification by the presence of epithelialized tunnels (Navrazhina et al., 2021) illustrated no significant difference between the number of SFRP1+ cells, but a decrease in the number of SFRP2+ cells and CXCL12+ cells in the absence of epithelialized tunnels compared to the presence of epithelialized tunnels (Figure 3d-f).

**Figure 3:**
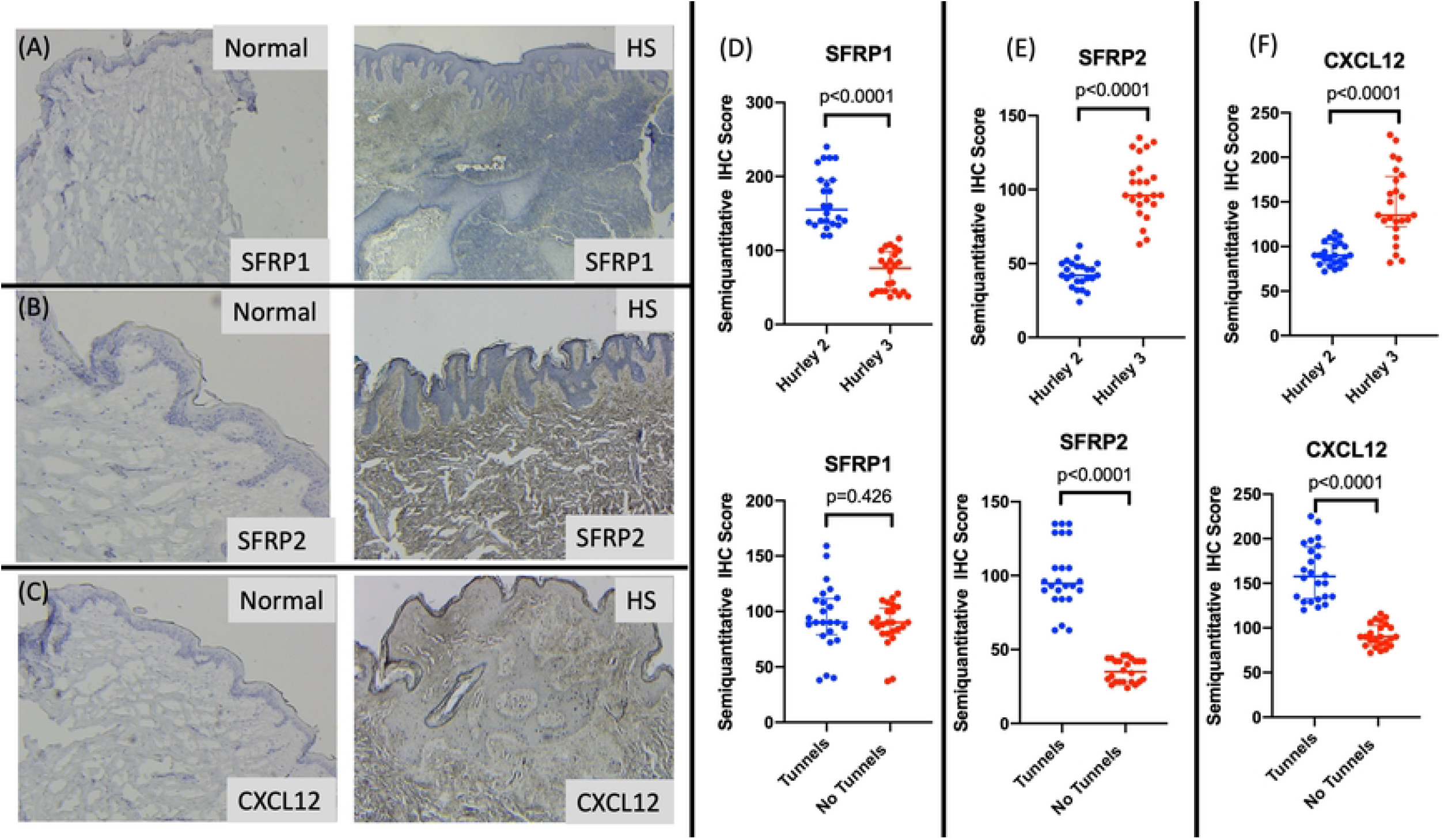
(A-C) Fibroblast subpopulations in HS differ based upon Hurley stage and presence of epithelialised tunnels. SFRP1 protein (D) is diffusely positive in the reticular dermis in Hurley stage 2 lesions but reduced in Hurley 3. No difference was seen in the presence of tunnels. SFRP2 protein (E) illustrates increased expression in Hurley stage 3 disease and in the presence of epithelialised tunnels. CXCL12 protein expression (F) is increased in Hurley stage 3 disease and in the presence of epithelialized tunnels.

### HS Lesions with epithelialized tunnels demonstrate elevated signals of Epithelial-to-Mesenchymal Transition in comparison to HS lesions without tunnels and healthy control tissue

Immunohistochemical staining for signs of epithelial mesenchymal transition (EMT) identified increased staining of TWIST1 (Figure 4a), ZEB1 (Figure 4b), Snail/Slug (Figure 4c) and N-Cadherin (Figure 4d) proteins in HS lesional tissue compared to site-matched healthy controls. The tips of elongated epithelial rete ridges in pseudo-psoriasiform hyperplasia demonstrate discrete clustering of positive staining to EMT-associated proteins including TWIST1 and ZEB1 (Figure 4a-b). Stratification of HS lesional tissue by the presence of epithelialized tunnels indicated a significant increase in the number of SLUG/SNAIL, ZEB1 and TWIST1 positive cells when compared with HS lesions without tunnels or healthy controls. (Figure 4e-g).

**Figure 4:**
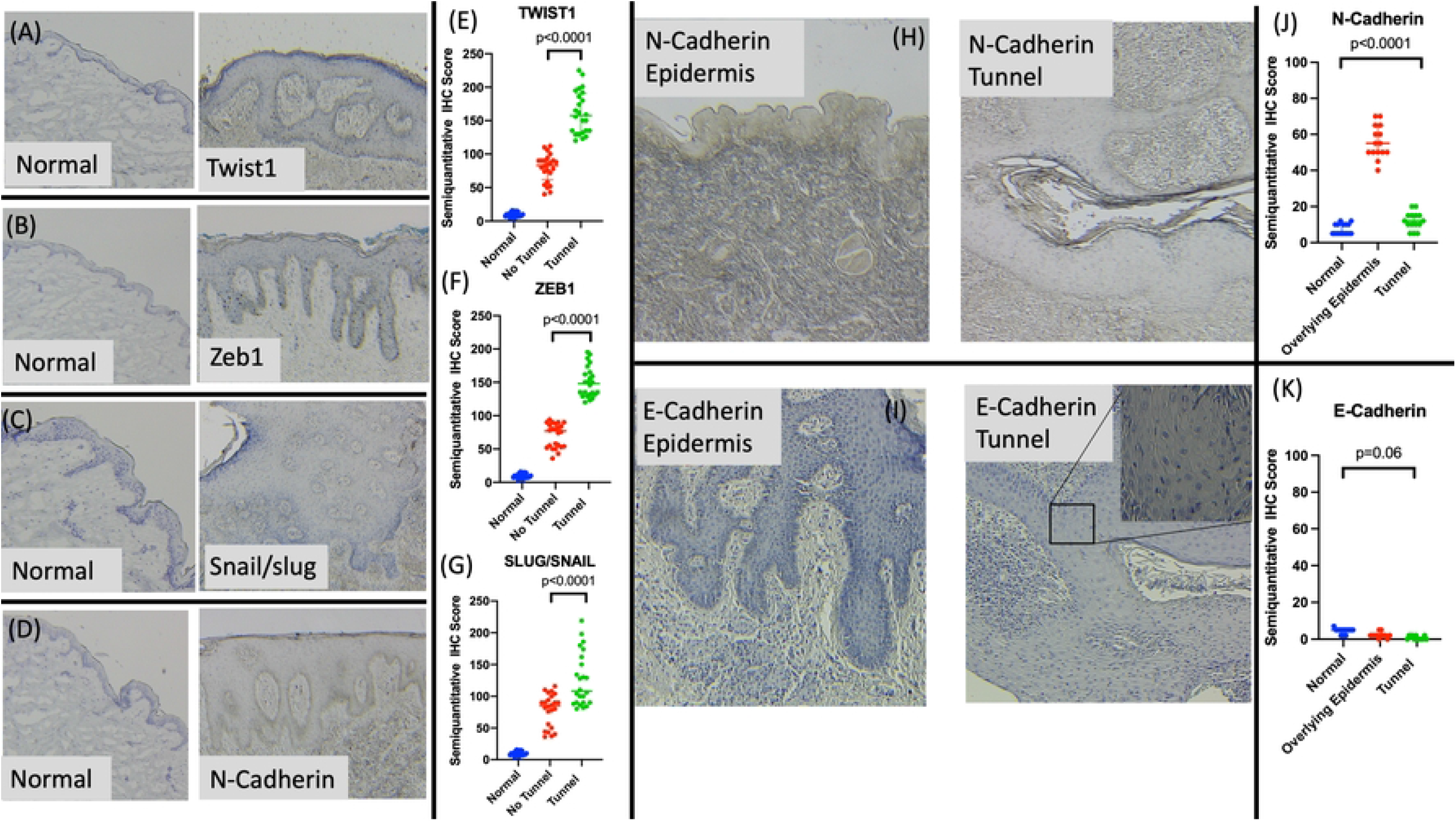
Epithelial Mesenchymal Transition Markers are significantly overexpressed in HS lesional tissue compared to site matched healthy controls (A-D). Protein expression of TWIST1 (E), ZEB1 (F) and SLUG/SNAIL (G) are significantly increased in HS lesional tissue with highest expression in lesions with epithelialized tunnels. Protein expression of N Cadherin (H) was significantly elevated in the overlying epidermis of lesions with tunnels but normal in the epithelial lining of the tunnel itself (J). E-Cadherin expression (I) showed no significant difference between sites (K).

Differences in intensity of N-Cadherin staining were observed between the overlying epidermal epithelium when compared with the epithelial lining of tunnels (Figure 4h-k). Significant increases in the number of N-Cadherin positive cells were seen in the overlying epidermal epithelium than epithelialised tunnels. The semiquantitative IHC staining in epithelialized tunnels was comparable to that of healthy controls (Figure 4j). Conversely, E-Cadherin protein expression is negligible in overlying epidermis but increased in tunnel epithelium compared to healthy controls, however this difference did not reach statistical significance (Figure 4k).

### Spleen Tyrosine Kinase Inhibition with Fostamatinib demonstrates significant reduction in Fibrosis, inflammatory and Epithelial-Mesenchymal Transition associated genes and pathways in Hidradenitis Suppurativa after 4 weeks of therapy

Spleen Tyrosine Kinase (SYK) inhibition with Fostamatinib at a dose of 100mg twice daily significantly modulates gene expression as measured by the Human Fibrosis V2.0 Nanostring gene expression assay. Fibroblast associated genes including *IL13, FGF21, COL3A1, VIM* and EMT associated genes including *SNAI2* and *ITGB1* were significantly downregulated at Week 4 in lesional HS tissue compared with baseline (Week 0) (Figure 5a,b). Pathway analysis demonstrated high levels of enrichment in the downregulation of gene ontology pathways associated with fibroblast proliferation, transforming growth factors beta production and connective tissue growth factor response as well as other inflammation associated pathways including regulation of chronic inflammatory response. (Figure 5c).

**Figure 5:**
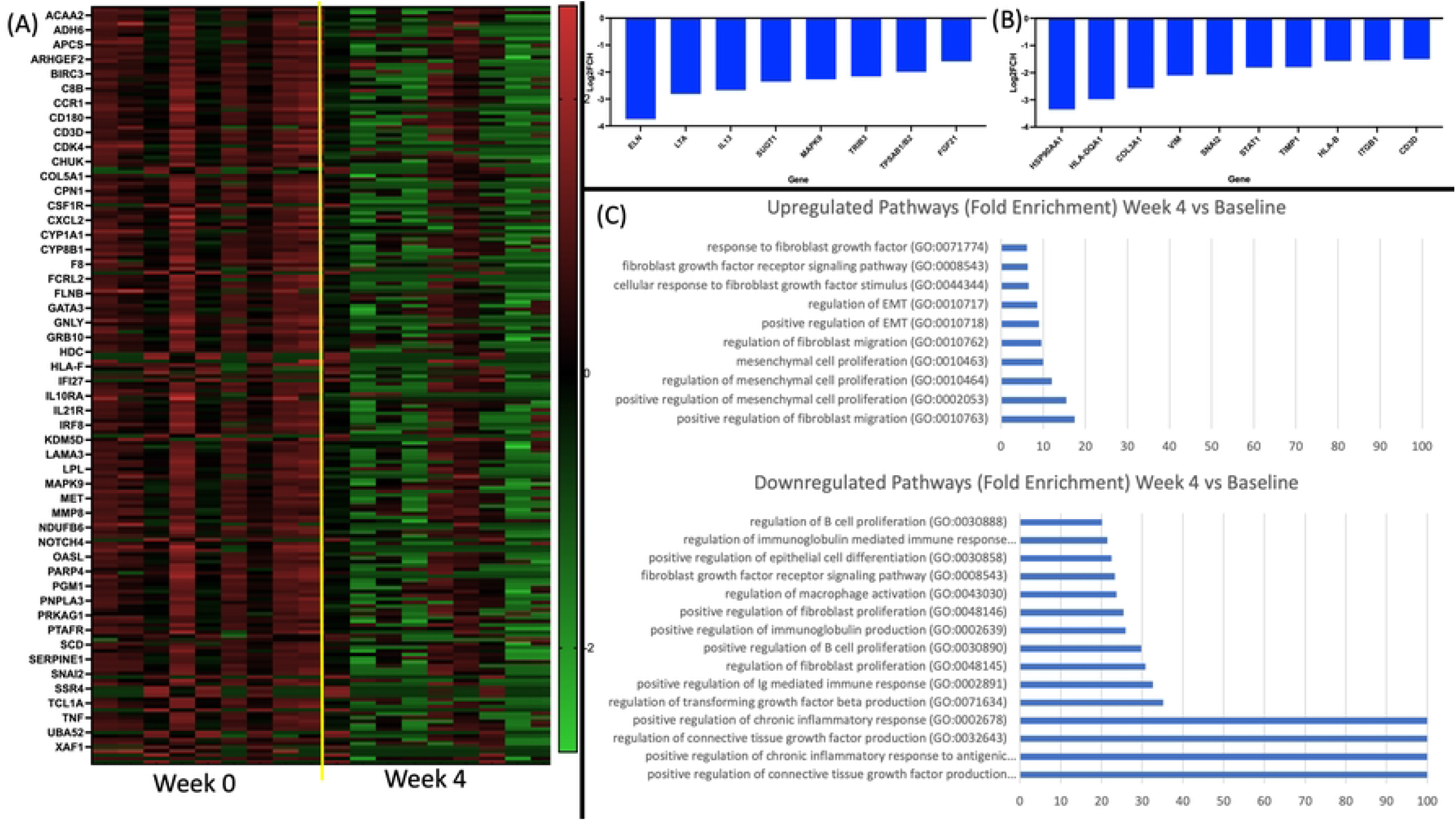
Therapy with 4 weeks of the oral spleen tyrosine kinase antagonist Fostamatinib downregulates gene expression relating to fibroblast activity and epithelial-mesenchymal transition as measured by Nanostring gene expression panel (A,B). Pathway analysis identified significant downregulation and enrichment of chronic inflammatory response, connective tissue growth factor and transforming growth factor beta production (C).

## DISCUSSION

This study presents novel gene expression and immunohistochemical data pertaining to the presence of fibroblast subpopulations and evidence of EMT in lesional tissue of HS. Fibroblast subpopulations in lesional tissue are significantly altered compared to non lesional tissue and healthy controls. Expansion of the SFRP2+ and CXCL12+ populations were identified with no significant alteration to the SFRP1+ population. Additionally, the severity of disease and presence of epithelialized tunnels is associated with these alterations to fibroblast subpopulations. Evidence of EMT was seen in HS lesional tissue with greater levels associated with epithelialised tunnels and Hurley stage 3 disease. SYK inhibition with fostamatinib for as little as 4 weeks resulted in reduced expression of fibroblast and EMT associated genes as measured by Nanostring gene expression assay. Pathway analysis of Nanostring data demonstrated enrichment of downregulated pathways pertaining to fibroblast proliferation and connective tissue growth factor production.

Previously published functional annotation of fibroblast subpopulations in normal human skin identifies SFRP2+ (Type A) cells as functionally associated with dermal cell and extracellular matrix homeostasis as well as inflammatory cell retention. Additionally, SFRP2+ cells are involved in the development of tertiary lymphoid structures (TLO). (Tabib et al., 2018). The significant expansion of SFRP2+ fibroblast subpopulations in HS lesional tissue would explain the development of TLOs (Wolk et al., 2020; Frew et al., 2020) in HS tissue. Additionally, gross histological finding sin HS include gross expansion of the dermal layer, fibrosis and retention of inflammatory cells, all associated with function of SFRP2+ fibroblasts.

CXCL12+ (Type B) cells contribute to immune surveillance and are associated with Th2 immune axis activation. This subset is particularly expanded in lesional tissue of atopic dermatitis. (He et al., 2019). SFRP1+ cells are described as being functionally responsible for ECM remodeling and include dermal papilla cells and fibroblasts of the dermal-hypodermal junction. The reduction in SFRP1+ cells in advanced Hurley stage 3 disease may be reflective of destruction of follicular units and decreases in the number of dermal papilla fibroblasts in tissue. Additionally, CXCL12+ cells were increased in severe disease and the presence of epithelialized tunnels suggesting an association with increased levels of inflammation (Navrazhina et al.,2021). Witte Handel et al (Witte Handel et al., 2018) has previously demonstrated fibroblasts as a major source of IL-1 and downstream inflammasome activation in HS, and our data supports the hypothesis that SFRP2+ and CXCL12+ fibroblasts are the subpopulations associated with this inflammatory signature. Further investigations with single cell RNA sequencing would be required to confirm this, as tissue culture results in preferential expansion of SFRP2+ cells (Tabib et al., 2021) which may lead to poor isolation of other fibroblast subsets.

Evidence of EMT has been reported in HS lesional tissue by Nelson and colleagues with reductions in E-cadherin in the epidermis (Nelson et al., 2019). This has been interpreted as a sign of sensitivity to mechanical stress (Danby et al., 2013), however our investigations in the epithelium of tunnels suggest that greater EMT signals are associated with epithelialised tunnels and may be indicative of a mechanism of tunnel development rather than a predisposing factor of disease initiation (Frew et al., 2019a). This is supported by the fact that non-lesional tissue did not demonstrate any significant differences in EMT protein expression compared to normal skin (Figure 4a-d), and that the EMT expression as measured by IHC was significantly elevated in biopsies with tunnels compared to those without (Figure 4e-g).

EMT can be induced in epithelial cells by inflammation and extracellular matrix alterations in both oncological and inflammatory settings (Suarez-Carmona., 2017; Wang et al., 2018). Additionally, the process of EMT stimulated the production of inflammatory cytokines including IL-1 (Suarez-Carmona., 2017). These results clearly demonstrate EMT signals in tissue of HS, however the precise causative cascade leading to the development of EMT in HS remains unclear. The use of Fostamatinib partially ameliorates the gene expression signature of EMT in HS tissue, presumably indirectly through its action upon B cells, plasma cells and monocyte/macrophages (Frew., 2022; Wang et al., 2021). Wang demonstrates direct alteration of fibroblast function with the use of fostamatinib in vitro (Wang et al., 2021) and therefore fostamatinib may have potential as an adjuvant treatment alongside other targeted therapies such as Janus Kinase Inhibitors or Monoclonal antibodies (Frew et al., 2021b), particularly in the setting of epithelialised tunnels and/or significant fibrosis.

The use of whole genome RNAseq has several disadvantages when examining discrete gene expression pertaining to cellular subsets (Whitley et al., 2016). Biases during cDNA library construction and sequence alignment, as well as read depth can alter transcript quantitation and reduce reproducibility between datasets. Single cell RNA sequencing is susceptible to drop out and significant noise bias due to the low amount of RNA analyzed. (Wu et al., 2018). Nanostring gene expression arrays addresses some of these disadvantages including no need for amplification, ability to identify low volume and degraded RNA and minimal background signal compared to whole tissue RNAseq. When RNAseq and Nanostring gene expression analyses have been compared in Psoriasis Vulgaris, increased accuracy for low level expressed transcripts was identified in Nanostring compared with whole genome RNAseq. (Krueger et al., 2019) In-silico deconvolution provides a computational alternative to scRNAseq which, when utilized with a high-quality reference frame can provide insights into cellular proportions in whole RNAseq data whilst avoiding the biases present in scRNAseq. The quality of the deconvolution data is predicated on the baseline RNAseq data. Whilst whole tissue RNAseq is most appropriate for biomarker discovery, for identification of a priori differential gene expression, the combination of in silico-deconvolution of whole tissue RNAseq and Nanostring gene expression (confirmed with immunohistochemistry) provides a robust methodology for investigation of pre-defined signatures of fibroblast subpopulations.

HS is in great need of novel therapies; however, our understanding of the disease pathogenesis and specific mechanistic action of novel therapies is incomplete. SYK is an important mediator of downstream signaling in monocytes, neutrophils and B cells but can also contribute to the reprogramming of fibroblasts. (Liu et al 2019) The positive clinical results in our recent open-label clinical trial of fostamatinib in HS (Jepsen et al., 2022) suggest that fostamatinib has a clinically significant impact on disease severity in HS. The analysis of gene expression data from the first 4 weeks of therapy suggests that a rapid alteration in chronic inflammatory and pro-fibrotic pathways is associated with this therapy and is associated with clinical response. Future analysis of this clinical trial data will enable deeper investigation into the precise mechanisms of SYK inhibition in HS at higher doses and further timepoints.

## MATERIALS AND METHODS

### Patients and Tissue Collection

This study was approved by the human research ethics committee of Sydney South West Area Health Service (Sydney Australia). A total of 20 participants underwent lesional and non lesional biopsies (6mm in diameter) based upon previously published definitions. All. Biopsies were immediately stored in RNA Later and frozen at -80C until processing. Standard wound care was undertaken for each of the biopsy sites. All patients were not on any active therapy at the time of biopsies. The clinical characteristics of participants are included in Supplementary Data.

### Skin Biopsy mRNA Quantification

Total RNA was extracted using the Qiagen RNeasy kit as per standard protocol (Qiagen, Valencia, CA, USA). RNA was sequenced using the Novaseq S4 (Illumina, San Diego, CA, USA) Differential expression was estimated with DESeq2 and VST to log_2_ transform the normalized counts. Differentially expressed genes were defined as fold change (FCH) of ≥ ¦1·5¦ and false discovery rate ≤ 0·05.

### Pathway Analysis

Enrichment analyses were completed using Protein ANalysis THrough Evolutionary Relationships (PANTHER) classification system (www.pantherdb.org). Analyses were performed using whole tissue RNAseq data (Figure 1) as well as for Nanostring Gene Expression Panels. Nanostring gene expression panels included Baseline (prior to fostamatinib administration) and Week 4 of fostamatinib therapy lesional tissue samples (Figure 5).

### CIBERSORTx

In silico deconvolution was undertaken using the CIBERSORTx web application ((https://cibersortx.stanford.edu/index.php) in order to estimate the abundance of fibroblast subpopulations in baseline lesional and non-lesional tissue using whole tissue RNAseq. The utilized fibroblast reference matrix was based upon previously published human skin scRNAseq data (GSE128169) with stratification into three major subtypes as defined by Ascension et al (https://doi.org/10.1016/j.jid.2020.11.028)

### Nanostring Analysis

Extracted RNA was analysed using the NCounter system (Nanostring) using the Human Fibrosis V2.0 gene panel. Raw data was processed using the Nanostring Nsolver (version 4.0.70) analysis software using quality control and normalization procedures derived from the NormqPCR R package as previously described (Bhattacharya et al., 2021). Differentially expressed genes (DEGs) were defined as >1.5 Log2Fold change with a false discovery rate<0.05.

### Immunohistochemistry

IHC staining was undertaken using primary antibodies listed in Supplementary Files. Stained slides underwent semiquantitative analysis using standardized, previously published methods. (Crowe et al., 2019) Differences between groups were analyzed using the Wilcoxon rank sum test for two variables and/or omnibus test using one-way ANOVA with adjustment for multiple comparisons was made using the Benjamini Hochberg procedure.

### Statistical Analysis

RNA sequencing statistical analysis was performed in R language (**www.r-project.org**; R Foundation, Vienna, Austria) using publicly available packages (**www.bioconductor.org**). Means of each group and differences between groups were estimated using least-square means. Hypothesis testing was undertaken using the general framework for linear mixed-effect models of the limma package. tissue was considered a fixed factors while random effect related to subjects was included. Unsupervised clustering was performed with the Euclidean distance and average agglomeration. RNAseq data *P*-values were corrected for multiplicity using Benjamini– Hochberg procedure.

### Data Availability Statement

Baseline RNA sequencing Data is publicly available using SRA Bioproject ID: PRJNA912704 Nanostring Data is publicly available using GEO Accession Number: GSE220454

## Acknowledgements

Nil

## Author Contributions

Akshay Flora: Investigation, Methodology, Visualization, Writing-Revision

Rebecca Jepsen: Investigation, Methodology, Writing-Revision

Emily Kozera: Methodology, Writing-Revision

Jane Woods: Methodoogy, Writing-Revision

Geoff Cains: Methodology, Writing-Revision

Michael Radzieta: Investigation Data Curation, Formal Analysis, Methodology, Writing-Revision

Matthew Malone: Investigation, Data Curation, Methodology, Writing-Revision

John Frew: Conceptualization, Data Curation, Formal Analysis, Investigation, Methodology, Software, Visualization, Writing-Original Draft Preparation

## Supplementary Figures

Supplementary Figure 1: Heatmap of whole tissue RNAseq in lesional versus non-lesional tissue in individuals with Hidradenitis Suppurativa.

Supplementary Figure 2: Individual patient cell propotion data from CibersortX in-silico deconvolution examining fibroblast sub-populations in lesional and non-lesional tissue of Hidradenitis Suppurativa.

Supplementary Table 1: Patient Demographics and Disease Characteristics

Supplementary Table 2: Differentially Expression Genes in whole tissue RNAseq

Supplementary Table 3: Immunohistochemical Antibodies used in this study

Supplementary Table 4: Differentially Expressed Genes in Nanostring Gene Expression Panel.

